# Neural network of superiority illusion predicts the level of dopamine in striatum

**DOI:** 10.1101/2022.02.01.478593

**Authors:** Noriaki Yahata, Ayako Isato, Yasuyuki Kimura, Keita Yokokawa, Ming-Rong Zhang, Hiroshi Ito, Tetsuya Suhara, Makoto Higuchi, Makiko Yamada

## Abstract

In evaluating the personality attributes and performance of the self, people are inclined to view themselves superior to others, a phenomenon known as superiority illusion (SI). This illusive outlook pervades people’s thoughts, creating hope for the future and promoting mental health. Although a specific cortico-striatal functional connectivity (FC) under dopaminergic modulation was previously shown to be implicated in SI, the underlying whole-brain mechanisms have remained unclarified. Herein, to reveal the neural network subserving individual’s SI, we conducted a data-driven, machine-learning investigation to explore the resting-state FC network across the whole brain. Using the locally-acquired resting-state functional magnetic resonance imaging data (*n* = 123), we identified a set of 15 FCs most informative in classifying individuals with higher-versus lower-than-average levels of SI in evaluating positive trait words (area under the curve [AUC] = 0.81). Among the 15 FCs, the contribution level to the classification was 11% by the previously-highlighted cortico-striatal FC alone, but 60% by the encompassing cortico-limbico-striatal network cluster. A newly-identified, cortico-thalamic FC and another FC cluster also demonstrated substantial contribution. The classification accuracy was generalized into an independent cohort (*n* = 36; AUC = 0.73). Importantly, using the same set of 15 FCs, we achieved prediction on an individual’s level of striatal dopamine D_2_ receptor availability (Pearson correlation, *r* = 0.46, *P* = 0.005). This is the first successful identification of the whole-brain neural network that simultaneously predicts the behavioral manifestation and molecular underpinning of an essential psychological process that promotes well-being and mental health.

**Significance Statement:** Superiority illusion (SI) is a basic self-referential framework that pervades people’s thoughts and promotes well-being and mental health. An aberrant form of SI has been reported in psychiatric conditions such as depression. Our hypothesis-free, data-driven investigation revealed the spatially-distributed neural network that for the first time achieved prediction on an individual’s levels of SI and the striatal dopaminergic transmission simultaneously. In principle, this multiple-biological-layer framework can be applicable to any behavioral trait to establish a link with its underlying neural network and neurochemical properties, which could quantitatively present the relation of its aberrant form with the pathophysiology of neuropsychiatric disorders. Future clinical research may aid in deriving a diagnostic biomarker for examining the related behavioral and neurochemical characteristics within individuals.

## Introduction

Personality and social psychology research indicate that people naturally embrace positive views in self-perception (1). The views are not just positive but often deviate from reality even among normal people, providing illusive, self-serving cognition about one’s personality, attributes, and performance (1, 2). One such representative form was conceptualized nearly four decades ago as superiority illusion (SI; also known as “above-average effect”), which refers to people’s inclination toward evaluating themselves as being superior to others (3–5). For example, in comparison to the average peer, most people find desirable traits (such as honest and sociable) to be more self-descriptive, whereas undesirable traits (boring and unpopular) less relevant (6, 7). Such a view is clearly illusive because, assuming a normal distribution, most people cannot be better than average. However, inflated positivity, or positive illusion, pervades people’s thoughts, affecting multiple facets of human cognitive, affective, and social functions, in a way that brings hope for the future, thereby promoting mental health (1). In fact, the diminished positive illusion has been reported in individuals with depressive symptoms who exhibit unrealistically negative predictions for future life events (8).

Previously, two neuroimaging studies provided clues to the neural mechanisms underlying SI (7, 9). A task-based functional magnetic resonance imaging (fMRI) study investigated the engagement of cerebral regions in SI while participants performed a self-evaluation task to determine the self-descriptiveness of verbal stimuli depicting positive and negative personality traits (7). The study focused on activities in seven specific regions of interest known to be implicated in self-reference, availability heuristics, and valence and emotional processings (7). The analysis revealed cerebral activities susceptible to the SI judgments in the medial prefrontal cortex (MPFC; Brodmann area [BA] 9/10), the dorsal anterior cingulate cortex (dACC; BA24), the posterior cingulate cortex (PCC; BA23), and the orbitofrontal cortex (OFC; BA 11/47) (7). Importantly, the individuals with a minimal level of SI exhibited enhanced activities in the OFC and dACC, suggesting their inhibitory role in the heuristic search process that facilitated SI judgments (7).

We further investigated the molecular underpinnings of SI using positron emission tomography (PET) and explored their relationship with the frontostriatal FC as measured by resting-state fMRI (rsfMRI) (9). Resting-state functional connectivity (RSFC) refers to the temporal correlation of low-frequency spontaneous activities of spatially distributed regions, which has been shown to reflect the history of activation and learning in the brain (10, 11). Recently, RSFC has been used to probe the functional integrity of the brain that promotes various cognitive, psychological, and pathological processes (11, 12). Our previous study was motivated by the fact that the medial frontal regions such as the dACC project to the striatum to compose the frontostriatal circuits mediating motivation and response selection in humans (13, 14). Since the striatum receives massive dopaminergic projections from the substantia nigra and the ventral tegmental area, it was hypothesized that the level of dopaminergic transmission could affect individual differences in SI through the mediation of the frontostriatal FC. Using a radioligand [^11^C]racropride with high affinity to the dopamine (DA) D_2_ receptor (D_2_R), we identified that the availability of striatal DA D_2_R was related to the individual’s level of SI, which was mediated by the frontostriatal FC between the sensorimotor striatum (SMST) and the dACC (9). The results suggested that DA acted on the striatal DA D_2_R inhibiting the SMST–dACC connectivity, thereby leading to SI enhancement. This finding was consistent with the first task-based fMRI study in which the dACC appeared to play an inhibitory role in generating SI (7). Collectively, the two previous studies demonstrated the pivotal role of the frontostriatal circuit in SI, for which DA neurotransmission acts as a controller of their activities, thereby affecting behavioral manifestations of SI (7, 9).

However, the neural underpinnings of SI may be more complex than considered by the previous region-of-interest-based investigations (7, 9). The SI necessitates evaluation of self-value against others warranting concomitant involvement of the distributed functional network of the brain that enables self-reference, social comparison, language processing, autobiographic memory, and so forth (7, 9, 15, 16). Furthermore, cerebral activities during these processing were shown to be differential based on affective valence of the content of evaluation (17–19). These results encouraged us to investigate the valence-specific neural manifestation of SI across the whole brain. Recently, the application of feature extraction techniques to a large data set of rsfMRI has propelled data-driven, hypothesis-free exploration of the RSFC, which is critically involved in behavioral and pathological attributes of interest (12). Importantly, this provides a common ground to quantitatively delineate the brain state pertaining to a particular attribute, and then investigates the relationship among multiple interrelated attributes. For example, we previously developed a machine-learning algorithm to identify the set of inter-regional FCs that were most relevant to the classification of patients with various psychiatric disorders and their normal controls (20–22). A metric derived from the set of machine-learning selected FCs was used to quantitatively predict the liability to a disorder of interest. Likewise, in the present study, an individual’s propensity to SI may be identifiable via an FC-based measure; consequently, its behavioral and neurochemical manifestations could be quantitatively interrelated.

Here, motivated by the arguments above, we present an RSFC-based investigation of the brain that is critically involved in an individual’s propensity to the SI. We applied our previously developed machine-learning technique to the rsMRI dataset (*n* = 123), and extracted the set of FCs that were informative in classifying populations with high and low propensity to SI. Upon confirming the reliability of the classification in an independent dataset (*n* = 36), we further attempted to predict individual levels of striatal DA D_2_R availability by using the properties of the same set of FCs. The outcome of the present study is the whole-brain RSFC-based representation of the SI that simultaneously predicts the individual behavioral level and the underlying neurochemical property.

## Results

### Behavioral Results

Two groups of healthy adults, the discovery cohort (*n* = 123; 31 females) and validation cohort (*n* = 36; all males) were established in the present study. See Table 1 for the detailed description on the demographic information in each cohort. Between the two cohorts, there were significant differences in age (two-sample *t*-test, *P* < 10^−7^) and sex composition (chi-square test, χ^2^ = 11.3, *P* = 0.001), which were considered in the feature selection procedure (Materials and Methods). The individual’s propensity to SI was measured separately for the positive and negative trait words in each cohort (Fig. 1 and Table 1). For the positive trait words, the mean and the standard deviation (SD) of the SI measurements (pSI) was 0.16 ± 0.20 and 0.19 ± 0.22 in the discovery and validation cohort, respectively. Adopting the mean value of the discovery cohort as a common threshold (pSI_mean_ = 0.16), each cohort was split into two subgroups, pSI_L_ and pSI_H_, wherein the participants had pSI values lower or higher than the pSI_mean_, respectively (Fig. 1*A* and Table 1). We note that adopting the threshold this way was necessary in the validation cohort in order to evaluate the generalization capability of the classifier constructed in the discovery cohort. Likewise, for the negative trait words, the mean ± SD of the SI measurements (nSI) was 0.13 ± 0.21 and 0.11 ± 0.21 in the discovery and validation cohort, respectively. Two groups, nSI_L_ and nSI_H_, were formed by dividing each cohort at the threshold of nSI_mean_ = 0.13 (Fig. 1*B*). In the discovery cohort, the mean age of the SI_H_ subgroup tended to be higher than that of the SI_L_ subgroup for both positive (*P* = 0.07) and negative (*P* = 0.06) trait words (Table 1). This trend in the age difference was considered in the subsequent feature selection procedure (Materials and Methods).

**Table 1.** Demographic information and the superiority illusion (SI) measurement for (*A*) positive and (*B*) negative trait words in the discovery and validation cohorts. \**P*-values were determined by a Wilcoxon rank sum test for SI, age, and BDI, and by a χ^2^ test for sex. †BDI (Beck Depression Inventory) measurements were available only for the discovery cohort.

| A. Positive trait words |  |  |  |  |  |  |  |  |
| --- | --- | --- | --- | --- | --- | --- | --- | --- |
| Item | Discovery cohort |  |  |  | Validation cohort |  |  |  |
|  | All | Subgroup |  | *P* * | All | Subgroup |  | *P* * |
|  |  | pSI<sub>L</sub> | pSI<sub>H</sub> |  |  | pSI<sub>L</sub> | pSI<sub>H</sub> |  |
| N | 123 | 64 | 59 | – | 36 | 14 | 22 | – |
| pSI | 0.16±0.20 | 0.01±0.11 | 0.33±0.13 | < 0.001 | 0.19±0.22 | –0.02±0.17 | 0.32±0.14 | < 0.001 |
| Age (y) | 31.4±13.6 | 29.3±11.4 | 33.5±15.4 | 0.07 | 23.3±4.4 | 22.9±3.5 | 23.6±4.9 | 0.84 |
| Sex (M/F) | 92 / 31 | 48 / 16 | 44 / 15 | 0.96 | 36 / 0 | 14 / 0 | 22 / 0 | – |
| BDI† | 5.5±4.9 | 6.4±5.7 | 4.5±3.6 | 0.13 | (n/a) | (n/a) | (n/a) | – |
| B. Negative trait words |  |  |  |  |  |  |  |  |
| Item | Discovery cohort |  |  |  | Validation cohort |  |  |  |
|  | All | Subgroup |  | *P* * | All | Subgroup |  | *P* * |
|  |  | nSI<sub>L</sub> | nSI<sub>H</sub> |  |  | nSI<sub>L</sub> | nSI<sub>H</sub> |  |
| N | 123 | 70 | 53 | – | 36 | 19 | 17 | – |
| nSI | 0.13±0.21 | –0.01±0.12 | 0.33±0.14 | < 0.001 | 0.11±0.21 | –0.03±0.19 | 0.26±0.10 | < 0.001 |
| Age (y) | 31.4±13.6 | 28.8±10.5 | 34.8±16.2 | 0.06 | 23.3±4.4 | 22.9±3.4 | 23.8±5.3 | 0.87 |
| Sex (M/F) | 92 / 31 | 53 / 17 | 39 / 14 | 0.79 | 36 / 0 | 19 / 0 | 17 / 0 | – |
| BDI† | 5.5±4.9 | 6.9±5.5 | 3.7±3.2 | 0.001 | (n/a) | (n/a) | (n/a) | – |

**Figure 1.**
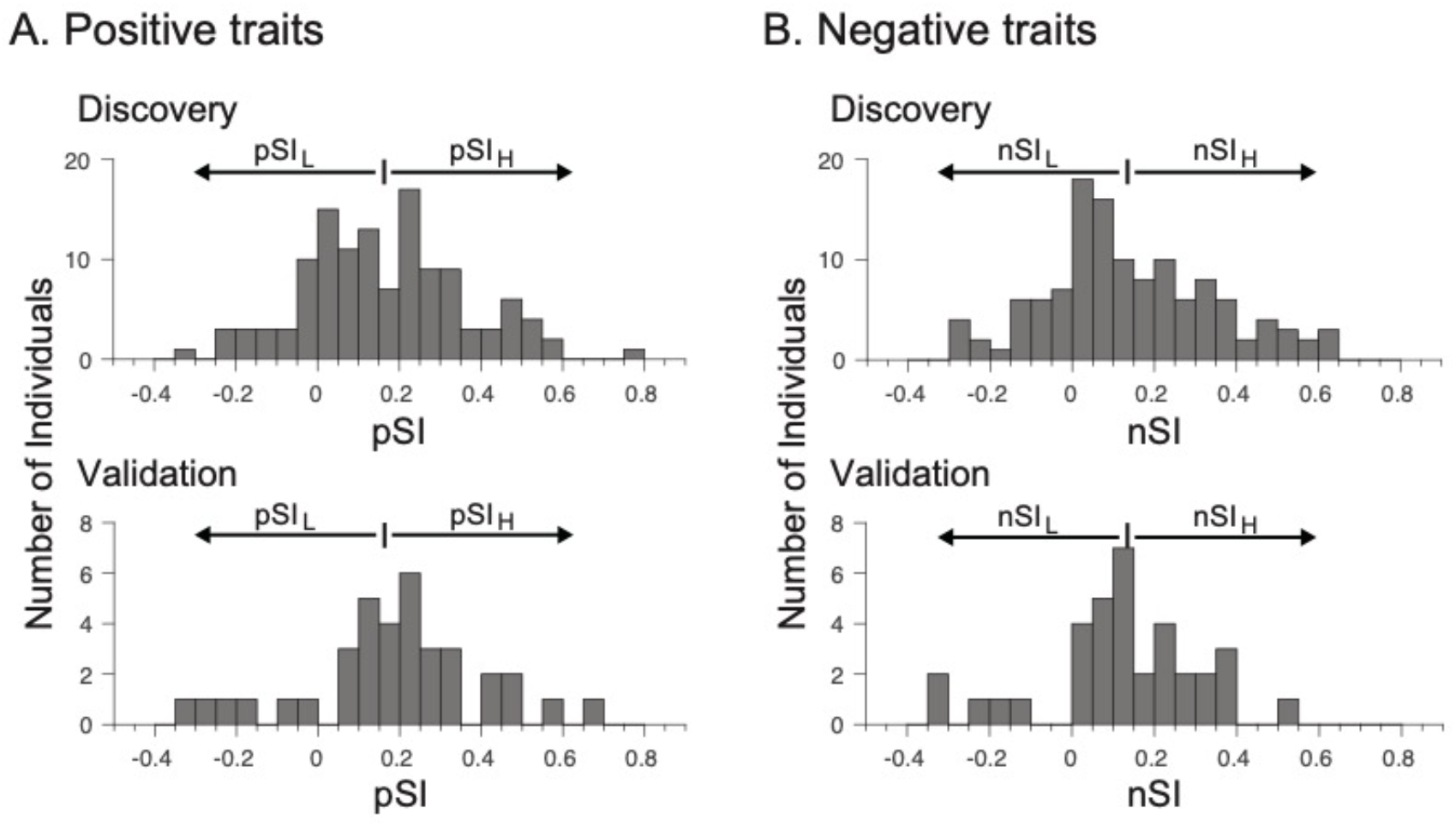
Distribution of the SI measurements for the (*A*) positive and (*B*) negative trait words in the discovery (top) and validation (bottom) cohorts. For each trait, the vertical line segment on the top of the histogram indicates the mean value of the superiority illusion (SI) measurement calculated in the discovery cohort. Two subgroups were formed within each cohort according to whether the individual has the SI value below (denoted by subscript L) or above (subscript H) the average value in the discovery cohort.

### Classification of populations above and below average SI

We sought the set of FCs informative to distinguish the SI_L_ and SI_H_ subgroups in the discovery cohort. Using the preprocessed rsfMRI datasets, we calculated the inter-regional correlation matrices which, for each participant, incorporated a total of 7,503 temporal Pearson correlation indices calculated among 123 regions of interest (ROIs) (Materials and Methods). The cascade of feature selection algorithms was applied to the pool of correlation matrices to identify the FCs that were most relevant to the distinction between the SI_L_ and SI_H_ subgroups while masking out the FCs affected by the covarying factors of the participants, such as age, sex, score of the Beck Depression Inventory (BDI), and head motion-related measurements (Materials and Methods).

For the positive trait words, the algorithms identified a total of 15 FCs distributed across the whole brain (Fig. 2 and Table 2). The weights determined in the feature selection procedure were then used to calculate the weighted linear summation (WLS) of the corresponding correlation indices of the selected 15 FCs (Materials and Methods). Specifically, we constructed a classifier in the form of a logistic function of the WLS by which an individual was classified into the pSI_L_ or pSI_H_ subgroups if the WLS was smaller or greater than zero, respectively. The leave-one-out cross validation (LOOCV) procedure revealed that the accuracy of the classification with respect to the actual subgroup identity was 73% (area under the curve [AUC] = 0.81; sensitivity = 80% and specificity = 67%) (Fig. 3), which was statistically significant (permutation test, *P* < 10^−4^). We confirmed that this classification scheme was generalized to an independent validation cohort with an accuracy of 72% (AUC = 0.73; sensitivity = 73% and specificity = 71%) (Fig. 3). Thus, we concluded that the WLS of the selected 15 FCs can be regarded as a neural network marker of the pSI that predicts one’s propensity to the SI for positive trait words.

**Table 2.** Properties of the 15 functional connectivities (FCs) used to classify the pSI_L_ and pSI_H_ subgroups. For each FC, the mean Pearson correlation index is calculated in each subgroup. Contribution was calculated as Weight × | *r*(pSI_L_) – *r*(pSI_H_) |, and the number in the parentheses indicates the relative fraction (%). Lat., laterality; BA, Brodmann area; Contrib., contribution; c., cortex; g., gyrus. *, see Figure 2.

| FC# | Terminal regions |  |  |  | Mean correlation |  | Weight | Contrib. (%) |
| --- | --- | --- | --- | --- | --- | --- | --- | --- |
| | Map ID* | Lat. | Name | BA | $r(\text{pSI}_L)$ | $r(\text{pSI}_H)$ | | |
| 1 | (5) | R | Thalamus | – | 0.08 | –0.03 | –9.47 | 1.04 |
|  | (15) | R | Middle frontal g. | 6 |  |  |  | (15.7) |
| 2 | (18) | R | Temporal fusiform c. (anterior) | 28 | –0.09 | 0.03 | 6.14 | 0.74 |
|  | (17) | L | Frontal pole | 10 |  |  |  | (11.2) |
| 3 | (1) | R | Putamen | – | 0.21 | 0.31 | 7.64 | 0.72 |
|  | (3) | R | Cingulate g. (anterior middle) | 24 |  |  |  | (10.9) |
| 4 | (12) | L | Cerebellum VI | – | –0.15 | –0.06 | 7.52 | 0.65 |
|  | (13) | L | Inferior frontal g. (pars triangularis) | 45 |  |  |  | (9.8) |
| 5 | (3) | L | Cingulate g. (anterior middle) | 24 | –0.07 | 0.03 | 6.24 | 0.63 |
|  | (6) | L | Occipital pole | 17 |  |  |  | (9.5) |
| 6 | (7) | R | Lateral occipital c. (superior) | 31 | 0.33 | 0.21 | –4.80 | 0.58 |
|  | (21) | R | Inferior temporal g. (temporooccipital) | 37 |  |  |  | (8.8) |
| 7 | (4) | L | Amygdala | 34 | –0.10 | 0.00 | 3.30 | 0.34 |
|  | (10) | R | Superior parietal lobule | 7 |  |  |  | (5.1) |
| 8 | (3) | L | Cingulate g. (anterior middle) | 24 | –0.21 | –0.32 | –3.03 | 0.34 |
|  | (7) | L | Lateral occipital c. (superior) | 19 |  |  |  | (5.1) |
| 9 | (8) | L | Cuneal c. | 18 | –0.10 | 0.00 | 2.65 | 0.26 |
|  | (7) | R | Lateral occipital c. (superior) | 31 |  |  |  | (3.9) |
| 10 | (19) | R | Parahippocampal g. (posterior) | 35 | –0.13 | –0.20 | –3.61 | 0.25 |
|  | (17) | R | Frontal pole | 10 |  |  |  | (3.8) |
| 11 | (11) | R | Cerebellum crus I | – | –0.01 | –0.12 | –2.41 | 0.25 |
|  | (4) | L | Amygdala | 34 |  |  |  | (3.8) |
| 12 | (16) | L | Frontal operculum c. | 13 | 0.04 | –0.05 | –2.56 | 0.22 |
|  | (20) | R | Middle temporal g. (temporooccipital) | 22 |  |  |  | (3.3) |
| 13 | (2) | R | Caudate | – | 0.12 | 0.04 | –2.86 | 0.22 |
|  | (4) | L | Amygdala | 34 |  |  |  | (3.3) |
| 14 | (21) | R | Inferior temporal g. (temporooccipital) | 37 | 0.15 | 0.04 | –1.89 | 0.21 |
|  | (14) | L | Inferior frontal g. (pars opercularis) | 44 |  |  |  | (3.2) |
| 15 | (9) | R | Supramarginal g. (anterior) | 40 | 0.11 | 0.01 | –1.88 | 0.17 |
|  | (10) | L | Superior parietal lobule | 7 |  |  |  | (2.6) |

**Figure 2.**
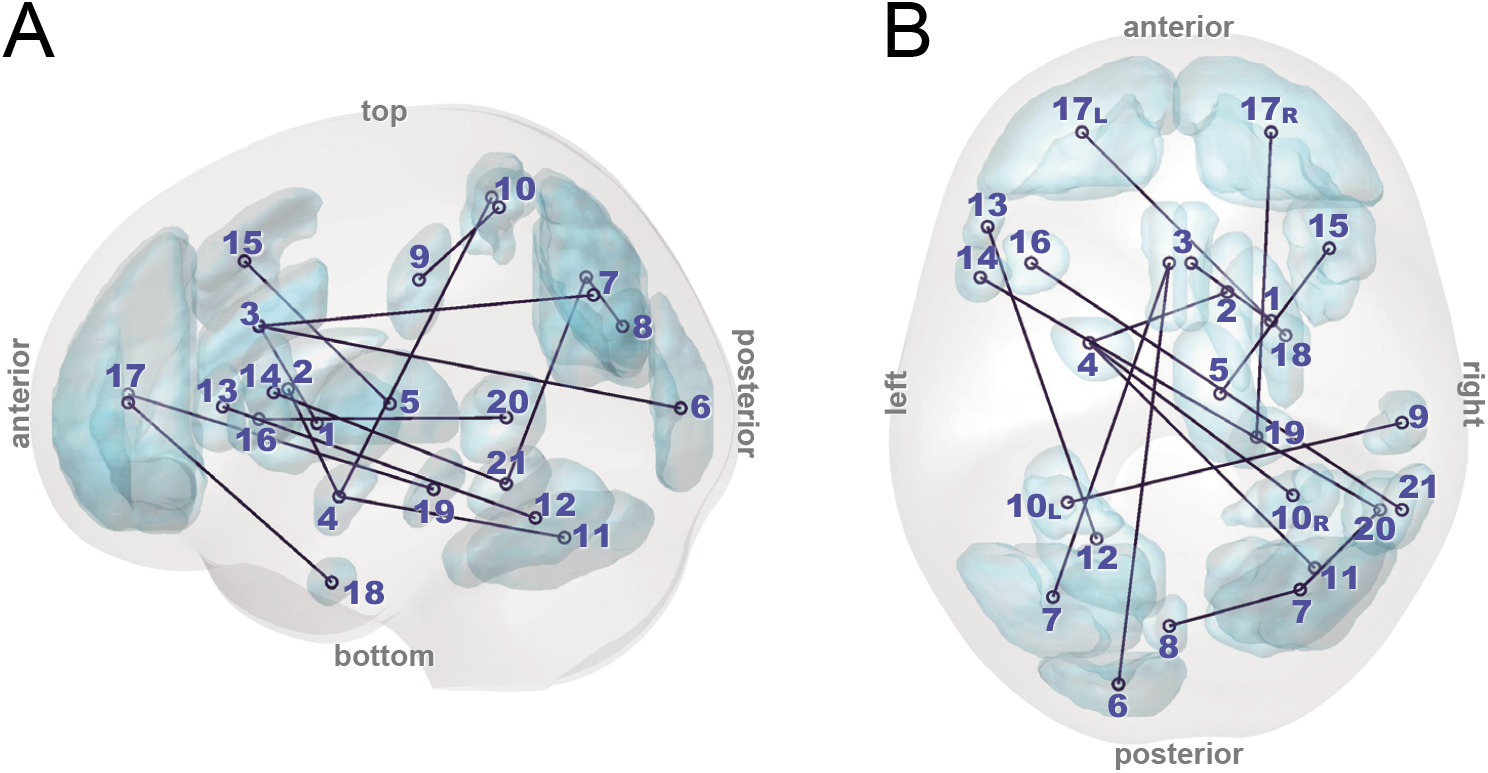
The 15 functional connectivities (FCs) used to classify the pSI_L_ and pSI_H_ subgroups as viewed from the (*A*) left side and (*B*) top of the brain. The ID numbers assigned to the terminal regions correspond to the Region ID in Table 2.

**Figure 3.**
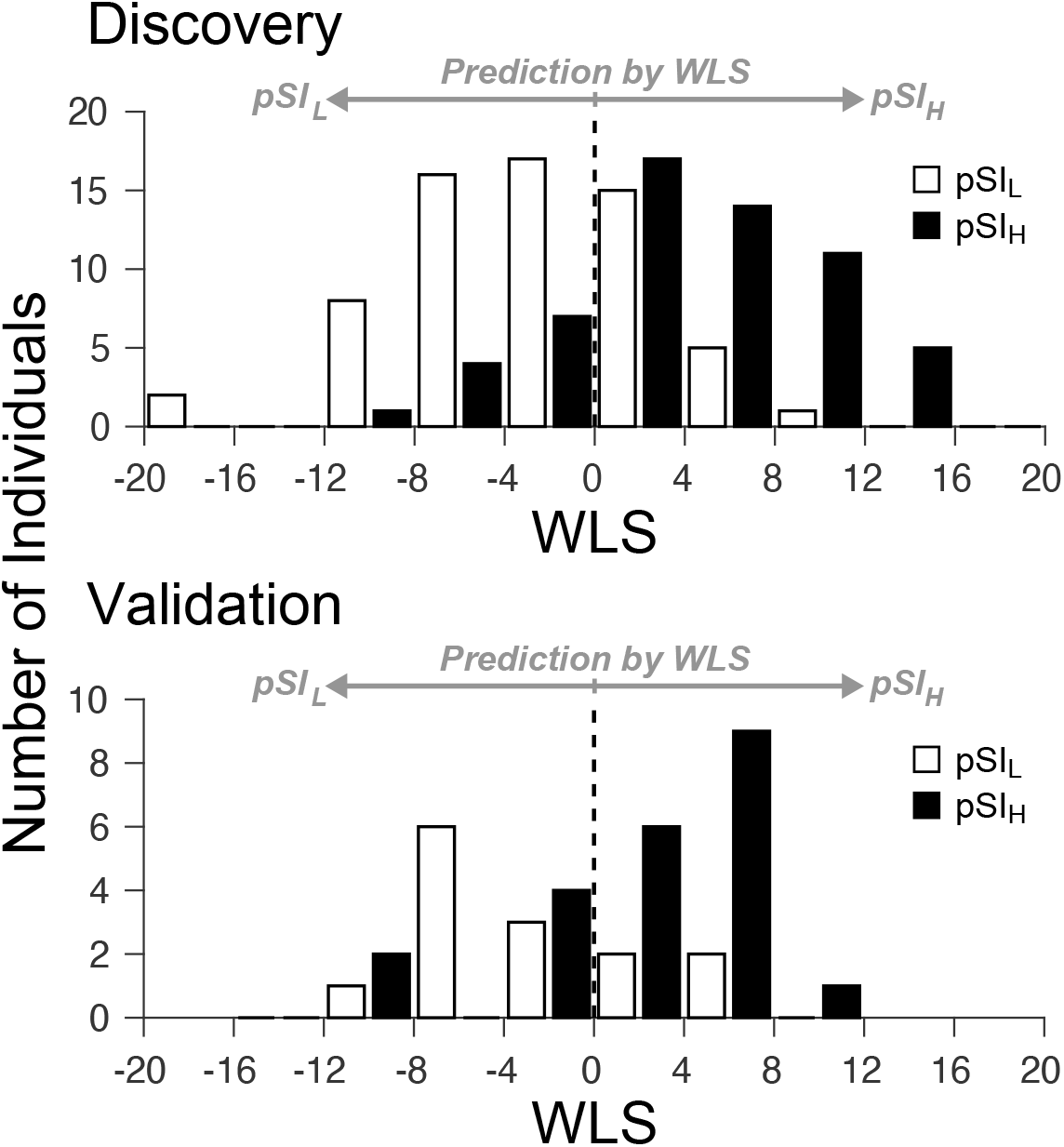
Distribution of weighted linear summation (WLS) of the 15 functional connectivities (FCs) selected by the machine learning algorithm (Materials and Methods) in the discovery cohort. The WLS smaller and greater than 0 is classified as pSI_L_ and pSI_H_ group, respectively. (*Top*) The number of individuals in the pSI_L_ (open bar) and pSI_H_ (filled bar) subgroups in the discovery cohort is shown as a histogram with the WLS width of 4. (*Bottom*) The WLS distribution of the validation cohort.

An identical analysis was repeated to evaluate the negative trait words. We were unable to identify the FCs that could classify the nSI_L_ and nSI_H_ subgroups (AUC = 0.41). We discuss the possible interpretation of this result in the Discussion, where a follow-up, preliminary analysis is described (see also Supplementary Information Text).

### Properties of the 15 FCs selected in the classifier for pSI subgroups

We focus on SI in the evaluation of the positive trait words. The 15 FCs selected in the pSI classifier were formed by 21 cortical and subcortical terminal regions that included two limbic structures (dACC and amygdala) connecting two striatal (putamen and caudate) and cortical regions, thereby forming a spatially distributed network across the brain (Fig. 4 and Table 2). In detail, we identified the following characteristics in the FCs. First, FC #3 is the connection between the putamen and the dACC, highlighted in our previous study (9). Its node, the dACC, acted as a hub, forming a cortico-limbico-striatal FC involving the occipital regions (FC #5 and #8). A similar pattern of connectivity was observed for FC #13, another limbico-striatal connection between the caudate and amygdala. Its node, the amygdala, acted as a hub, extending the connectivity to the parietal cortex (FC #7) and the cerebellum (FC #11). Second, FC #1 is a connection between the thalamus and the middle frontal gyrus, presenting as the most contributing FC to the classification of pSI_L_ and pSI_H_ subgroups (color-coded in yellow in Fig. 4). The contribution level to the classification was 16%, which was higher than that of the previously-highlighted FC #3 (11%) (see Table 2). The previous PET studies revealed the distribution of DA D_2_-like receptors in the human thalamus (23). FC #1 could therefore present as an alternative FC that bridges the site of DA transmission and the prefrontal cortex. Third, most FCs (8 out of 15) are cortico-cortical connections (FC #2, #4, #6, #9, #10, #12, #14, and #15) that incorporated various frontal and temporal regions not directly related to the striatum. Overall, two large network clusters emerged from these FCs (orange and green nodes in Fig. 4), each incorporating distinct limbico-striatal FC, i.e., one involving FC #3 and another involving FC #13. Altogether, these results underscored the importance of exploring whole-brain FCs to attain a comprehensive picture of the neural mechanisms subserving SI.

**Figure 4.**
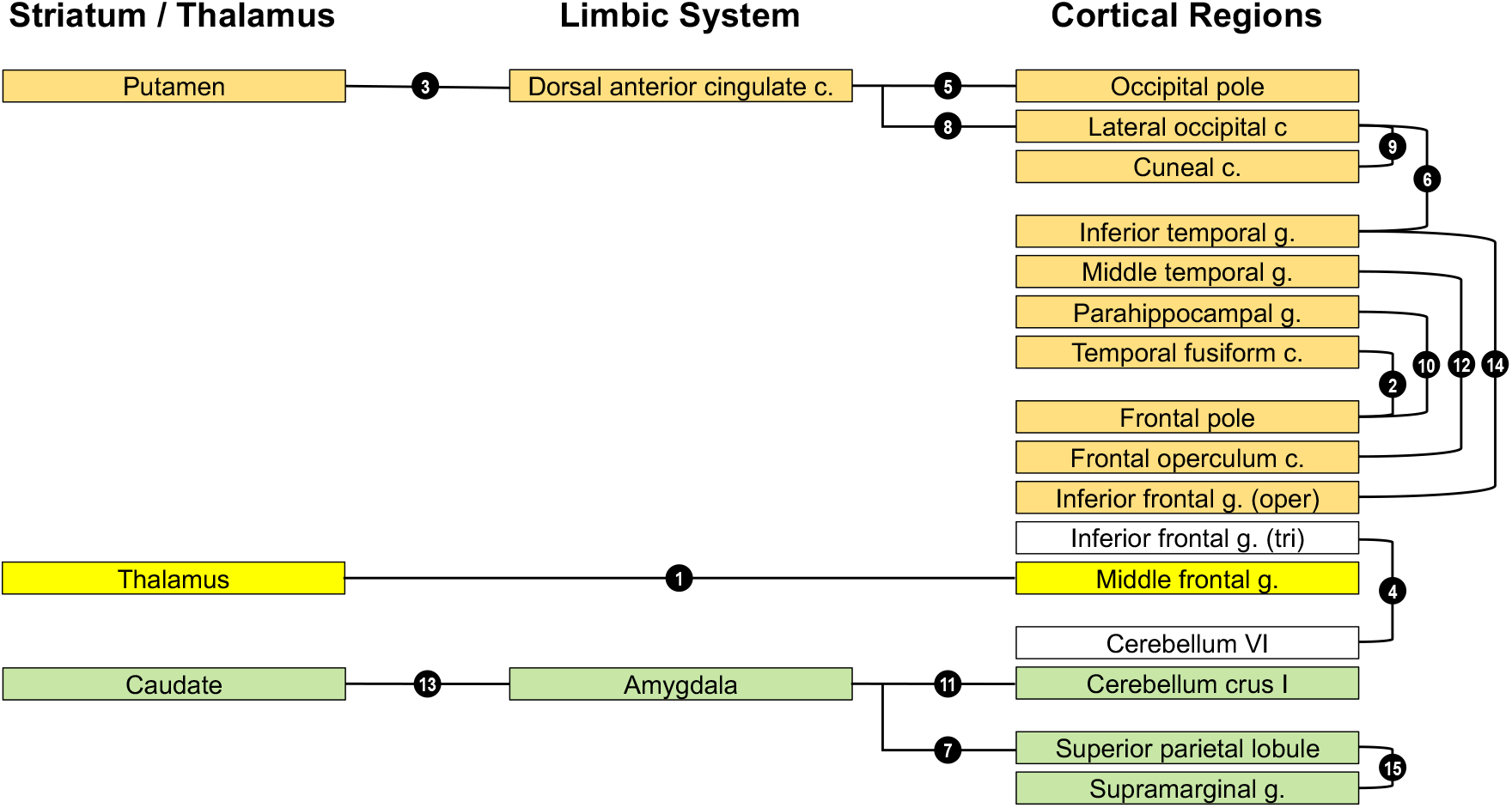
A schematic diagram depicting the mutual relationship among the 15 functional connectivities (FCs) used in the classification of the pSI subgroups. The number in the filled circle corresponds to the ID number in Table 2. The 15 FCs included two clusters of FCs, color-coded in orange and green, and their total contribution levels to the classification were 60% and 15%, respectively (see Table 2). The former cluster included the previously-highlighted cortico-striatal FC #3 (9). The FC #1 is an isolated cortico-thalamic FC, color-coded in yellow, which exhibited the highest contribution level as a single FC (11%).

We confirmed the significance of the selected 15 FCs in the classification of the two pSI subgroups based on their weights assigned in the feature selection procedure as follows: First, the cumulative absolute weights across the LOOCV indicated that these 15 FCs constituted an important subset of the 47 (out of 7,503) FCs that were selected at least once throughout the LOOCV procedure (Fig. S1). Second, for each of the 15 FCs, the set of weights assigned in the LOOCV procedure was significantly non-zero (one-sample *t*-test, *P* < 0.019 adjusted for false discovery rate; see Materials and Methods). These results demonstrate the significance of the 15 FCs in the classification of the pSI subgroups.

### Prediction of individual DA D2R availability by the 15 FCs in the pSI classifier

We investigated the relationship between the propensity to the SI for positive trait words and striatal DA D_2_R availability, as measured by the non-displaceable binding potential (BP_ND_) of a PET radioligand [^11^C]raclopride. As a target ROI, we focused on the left sensorimotor striatum (SMST) whose FC with the dACC was previously shown to be significantly correlated with BP_ND_ (7). In the present analysis, the BP_ND_ in the left SMST negatively correlated with individual pSI’s (Figure S2; *r* = –0.43, *P* = 0.008). In the group level, the mean ± SD of the BP_ND_ was 2.44 ± 0.23 and 2.29 ± 0.25 for the pSI_L_ and pSI_H_ subgroups, respectively. We observed that the former was significantly higher than the latter (*P* = 0.034, Wilcoxon rank sum test) (Figure 5A). This replicates our previous finding (7) using a larger sample.

**Figure 5.**
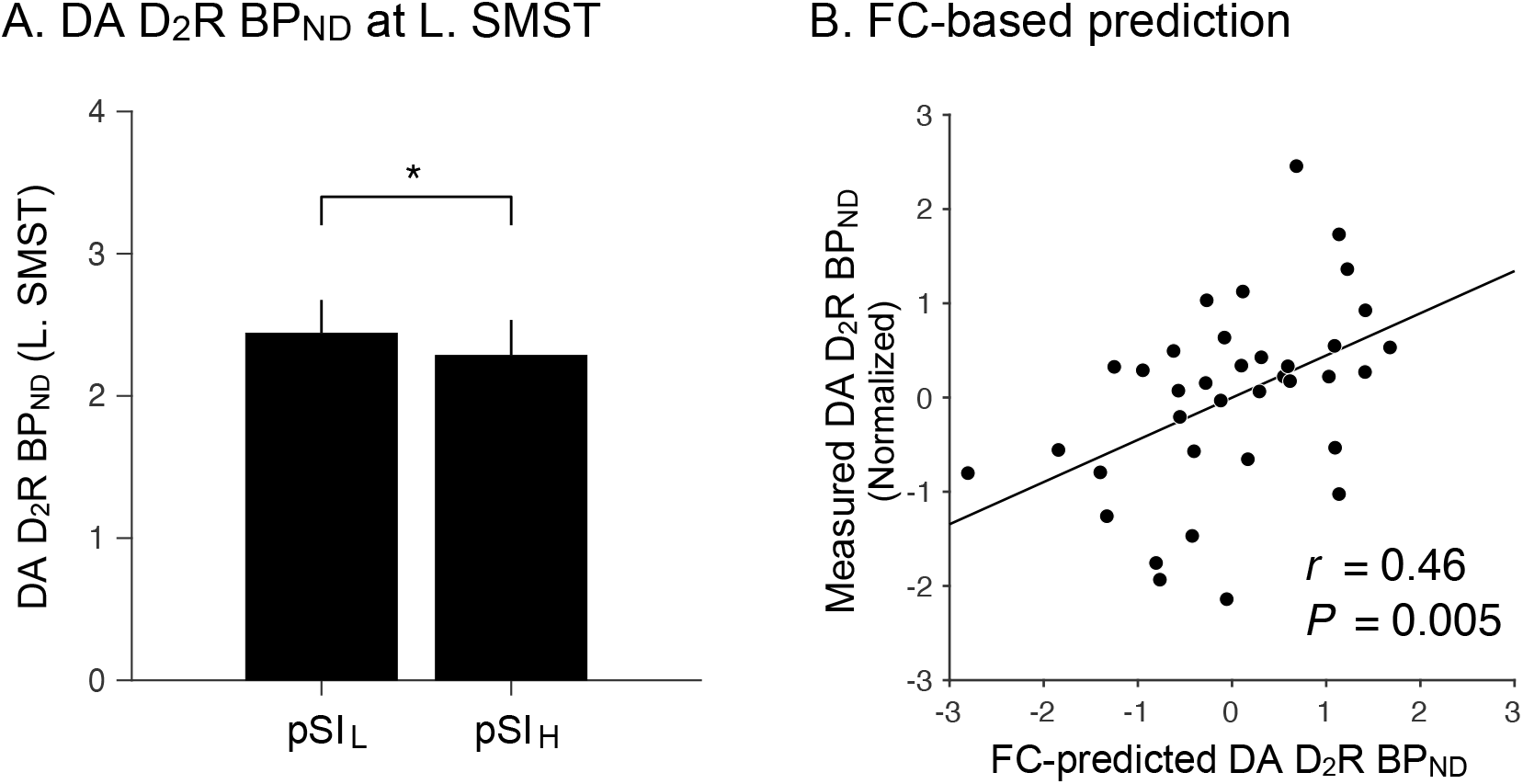
(*A*) The DA D_2_R availability in the left sensorimotor striatum (SMST) of the validation cohort (*n* = 36) as measured by non-displaceable binding potential (BP_ND_) of a radioligand [^11^C]raclopride. The pSI_L_ subgroup exhibited significantly higher DA D_2_ BP_ND_ than the pSI_H_ subgroup (*P* = 0.034, Wilcoxon rank sum test). (*B*) Prediction of the DA D_2_ BP_ND_ using the correlation indices of the 15 FCs. The actual and the predicted measurements were correlated significantly (*r* = 0.46, *P* = 0.005).

Furthermore, we examined whether the set of FCs for the pSI classifier could be used to predict an individual’s DA D_2_R availability in the striatum (Figure 5B). In the LOOCV framework, a linear regression model involving the correlation indices of the 15 FCs was fitted to the BP_ND_ measurements in the left SMST; thus, the model with the derived parameters could predict the BP_ND_ of the held-out individual. We observed that the measured and predicted BP_ND_ measurements correlated significantly (*r* = 0.46, *P* = 0.005) (Figure 5B). The robustness of the prediction was further confirmed by the permutation test (*P* = 0.003, 10,000 repetitions). Thus, we concluded that the set of 15 FCs was the neural network manifestation of the SI that interfaced its behavioral and molecular counterparts simultaneously.

## Discussion

Here, we presented an rsMRI-based investigation to explore the RSFC of the whole brain that underlies an individual’s propensity to SI. The machine learning-based feature selection algorithms identified the set of 15 FCs that were most relevant in SI for positive traits. The WLS of their respective correlation indices reliably predicted an individual level of pSI, that is, below or above the population average. The robustness of the classification was confirmed using an independent validation cohort. Using the same set of 15 FCs, we succeeded in predicting the individual’s level of DA D_2_R availability in the SMST. To the best of our knowledge, this is the first successful identification of RSFC that neurally subserves a specific mental process and that allows the simultaneous prediction of behavioral and molecular manifestations within the same individual.

The SI entails self-referential and social comparative processing to enable evaluation of self-value against others, suggesting a spatially distributed functional network of the brain that underlies the SI. Previous task-based fMRI studies on the self-evaluation of personality traits highlighted the central role of the midline structures, such as the MPFC, ventral and dorsal ACC, and PCC, whereas other regions in the prefrontal, temporal, and parietal cortices exhibited concomitant activities modulated by emotional and semantic attributes of the evaluated stimuli (7, 9, 15, 16). Social comparison has been mainly studied in competitive settings, and a recent meta-analysis on 59 extant studies has revealed that downward social comparison (i.e., being better than others) consistently activated the striatum and MPFC, while upward comparison (i.e., being worse than others) recruited the insula and dACC activation (24). The present study provides a comprehensive picture of the default functional relationships among the regions implicated in SI (Fig. 4). Specifically, in addition to replicating our previous finding of the putamen (SMST)–dACC connectivity as FC #3 (7), we demonstrated that this FC was a part of the large network cluster composed of frontal, temporal, and occipital regions, overall exhibiting 60% of the contribution level to the classification (orange nodes in Fig. 4). While it may be of future interest to find the specific roles for these regions in SI, our recent work has shown that resting-state functional networks in the frontal and temporal regions are associated with positive memory-specific recollection (25). Other FCs selected by the present algorithm included the cortico-thalamic connection (FC #1) and the cortico-limbico-striatal connections comprised of the caudate, amygdala, and a parietal region (FC #7 and #13). It is noteworthy that the dACC, striatum, and thalamus are major nodes of the salience network involved in self-regulation of cognition, behavior, and emotion (26). The current findings suggest that the cortico-striatal and cortico-thalamic connections, which are parts of the salience network, appear to be central to mechanisms of cognitive control associated with SI (7, 9). Both functional and structural abnormalities in the salience network have been observed in several psychiatric disorders, such as depression and schizophrenia (27). Thus, aberrations of control over SI may take a form of depressive realism in case of over-suppression, and a form of delusion of grandeur in case of insufficient suppression on the urge to expect positive self-view.

Although the present study focused on the SI, positive illusion can project as other forms of manifestation known as optimism bias and illusion of control (1). Optimism bias refers to an over- and under-estimation of the likelihood of experiencing positive and negative events, respectively (28). In a previous fMRI study, neural correlates were sought by contrasting the activities when the participants imagined future positive versus negative events (29). Enhanced activity was found in the amygdala, the rostral ACC, the caudate, and other regions known to be implicated in autobiographical memory retrieval and future projections, such as the inferior and medial frontal gyri, and middle temporal gyrus (30). The illusion of control refers to a phenomenon where one expects a success probability that is inappropriately higher than the objective probability would warrant (31). A previous fMRI study investigated subjective belief in control in an uncertain gambling setting (32). When comparing neural activities between the groups of participants with and without experience of the illusion of control, the former group exhibited increased activity in the nodes of the cortico-striatal network, including the nucleus accumbens and the right inferior frontal gyrus (32). Based on these previous findings, it is suggested that the three forms of positive illusion, the SI, optimism bias, and illusion of control, may have a common neural mechanism by sharing the nodes of connections in the striatum and prefrontal cortex.

The methodological framework to identify SI-related FCs worked successfully for positive trait words but not negative trait words. We speculate the possibility of inhomogeneity in the discovery cohort, that is, the presence of subgroups pertaining to the processing of negative trait words. Previous studies have suggested that individuals with high anxiety, such as social anxiety disorder, demonstrated altered patterns of neural activity during negative self-referential processing (33). Individuals with high and low levels of anxiety may thus exhibit differential involvement of RSFC in the manifestation of SI for negative trait words. In the feature selection process, the existence of heterogeneous subgroups in a single dataset makes identifying FCs that represent the entire group difficult. As an illustrative example, we previously employed a similar scheme for the investigation of major depressive disorder (MDD) (20). The scheme initially failed to identify FCs that distinguished groups of patients with MDD and normal controls to a meaningful accuracy (AUC = 0.62). However, considering MDD a generic label for a constellation of heterogeneous subtypes, we narrowed the scope of analysis into one major subtype of MDD, namely, the melancholic MDD characterized by anhedonia and lack of reactivity to pleasurable stimuli, and other symptoms (34). We then found that the extracted set of FCs accurately distinguished the melancholic patients with MDD from the control (AUC = 0.91); this classification scheme was generalized to an independent cohort (20). This underscores the fact that the efficacy of feature extraction depends on the homogeneity of the population. In the present study, the limited sample size of the dataset and incomplete demographic information did not permit a full investigation of the validity of the current argument. We present a follow-up analysis in the Supplementary Information Text. We attempted feature selection in a subsample of the discovery cohort (*n* = 95) in which individuals with high state anxiety were maximally excluded using the available information. The successful classification result (AUC = 0.78) in the subsample encourages a future in-depth study to confirm the validity of the current proposal.

We acknowledge the following limitations in this study. First, feature selection using MRI data is generally predestined to exploit nuisance variables (NVs) unique to a given sample data, and select features correlated with the NVs (21). This results in overfitting of the sample data, impairing its generalizability to independent data (35). NVs include both demographic factors and instrumental biases (21, 36). In the present study, we used the previously established technique (L_1_-SCCA algorithm) (21) to identify and mask out the features correlated with the demographic (age, sex, and BDI score) and measurement factors (head motion-related parameters). However, since the discovery data were acquired in a single protocol at a single site, it is possible that the feature extraction was biased owing to the particular settings in the discovery data, such as the choice of imaging apparatus and parameters. Future work should therefore incorporate multi-site MRI data and thereby confirm the reproducibility of the SI-related feature extraction by optimally factoring out instrumental biases by applying post-hoc analytic algorithms (e.g., (36)). Second, considering the limited sample size of the datasets, the reliability of the feature selection, the classification between the high/low SI subgroups, and the prediction of individual DA D_2_R availability should be further evaluated in a larger population. In addition, the use of various personality questionnaires and psychological instruments related to anxiety and depression could help clarify the mechanisms underlying SI for negative trait words. Third, since [^11^C]racropride is most sensitive in the striatum where the DA D_2_R density is high (~30 pmol/mL) (37), the present results do not explain the role of the less-dense (< 2.5 pmol/mL) (38), extrastriatal DA D_2_R in SI manifestation. Since the SI entails a multitude of cognitive and affective functions involving the prefrontal cortex and the limbic system, exploring the extrastriatal DA D_2_R function using other radioligands may complement the findings of the present study. As an intriguing case, a previous study used [^11^C]racropride and [^11^C]FLB 475 to measure the binding of the striatal and extrastriatal DA D_2_R, respectively, clarifying their distinct roles in different aspects of social desirability (39). A similar approach may help establish a more comprehensive picture of the molecular and neural mechanisms underlying SI.

In conclusion, we conducted a data-driven, machine-learning-based investigation and identified a set of 15 cortico-limbico-striatal, cortico-thalamic, and cortico-cortical FCs that were most informative in classifying two groups with pSI higher and lower than the group average. Using the same set of FCs, we were able to predict the individual levels of DA D_2_R in the SMST. Our study clarified how an individual’s neurochemical and neural network properties interact with each other to manifest SI-related behavior, an essential psychological process promoting well-being and mental health. We believe that the present methodological framework would be applicable to various other behavioral traits to substantiate our understanding of the etiology and pathophysiology of related neuropsychiatric disorders across multiple biological layers in the brain.

## Materials and Methods

### Participants

The present study was approved by the Ethics and Radiation Safety Committee of the National Institute of Radiological Sciences, National Institutes for Quantum and Radiological Science and Technology, Japan, in accordance with the ethical standards laid down in the 1964 Declaration of Helsinki and its later amendments. All participants provided written informed consent prior for participation in the study; all had normal or corrected-to-normal vision and had no history of neurologic or psychiatric disorders. No participant was taking any medications that could interfere with the interpretation of the results presented here. Two groups of participants, the discovery and validation cohorts, were established in the present study (Table 1). A total of 123 (age, mean ± SD = 31.4 ± 13.6; 31 females) and 36 healthy adults (age, 23.3 ± 4.4; all males) were included in the study for the discovery and validation cohort, respectively. The subset of the validation cohort (*n* = 24) was also used in our previous study (9). The Japanese version of the Beck Depression Inventory (BDI) (40, 41) was administered to the discovery cohort (mean ± SD = 5.5 ± 4.9). In the feature selection step (see below), the age, sex, and BDI scale were treated as NVs to mitigate their confounding effects in the feature selection and thereby improve the generalization capability of the derived classifier.

### The SI measurement

The procedure for SI measurement has been described in detail elsewhere (9). In brief, fifty-two socially desirable (positive) and undesirable (negative) trait words were selected from the previous literature (42) and translated into Japanese. Outside the scanner, participants were asked to rate how distant they were from the average peer on these personality traits using a visual analogue scale (ranging from 0 to 100 with an average of 50), yielding SI measurements. To derive the magnitude of the SI, the mean deviation from the average of 50 was calculated by reverse-scoring the ratings of negative traits to collapse with ratings of positive traits for each participant. Finally, the range of the SI measurements were linearly rescaled from [0,100] to [−1, 1].

### MRI data acquisition and analysis

All the participants underwent an MR scan with the imaging parameters and procedures summarized in Table S1. We utilized the CONN Toolbox (version 18b) (43) running with SPM12 (Wellcome Trust Centre for Neuroimaging, University College London, UK) software on MATLAB (R2018a, Mathworks, USA) to preprocess and denoise the raw data and to calculate the mean time course within the regions of interest. We used the default preprocessing pipeline that included realignment of the functional volumes to account for head motion, slice-timing correction, spatial normalization of the images to the Montreal Neurological Institute (MNI) space, and spatial smoothing using a Gaussian of full width at half maximum of 6 mm. To eliminate physiological and other noise of non-neuronal origins from the time course of blood oxygenation level dependent (BOLD) signal, subject-level denoising was performed using a regression model with the following confounds: (i) six motion parameters, their first-order derivatives, and their quadratic effects, (ii) five principal components in each of the white matter and cerebrospinal fluid (44), (iii) mean time course within the gray matter mask, and (iv) binary flags indicating the scan numbers where excessive frame-to-frame motion was detected. In (iv), the head motion was evaluated using the CONN’s built-in algorithm (ArtRepair (45)) with conservative thresholds (global signal *z*-value = 3, subject motion = 0.5 mm). The residual was then band-pass filtered in the range of [0.01, 0.1] Hz, yielding the BOLD time course for use in the subsequent calculation.

### Construction of composite atlas and calculation of interregional correlation matrices

In the present study, FCs were evaluated on an anatomical region basis using a composite atlas consisting of the Harvard-Oxford cortical and subcortical atlases (46) and the probabilistic cerebellar atlas (47) to cover the entire brain. The following two modifications were established based on this composite atlas. First, because previous studies on neural mechanisms of self-referential processing commonly used fine parcellation into the anterior cingulate cortex (ACC) (48), the corresponding area in the Harvard-Oxford atlas was parcellated into the three subregions, that is, the perigenual, anterior middle, and posterior middle portions of the ACC. The anterior middle portion corresponds to the dACC. Second, visual inspection of the functional images in the present datasets revealed that the lobules Crus II and VII through X of the cerebellum fell outside the field of view in some participants. Therefore, these subregions in the cerebellum were excluded from the analysis. After these two modifications, the final composite atlas contained a total of 123 regions (100 cortical, 14 subcortical, and 9 cerebellar regions). For each participant in the dataset, we calculated the mean time course of the BOLD signal in each region using the CONN toolbox following subject-level denoising. Finally, we calculated a pairwise interregional correlation matrix among 123 ROIs containing 7,503 (= 123 × 122 / 2) correlation indices.

### Feature Selection

Using the previously developed machine-learning algorithms (20–22), we identified the set of FCs by which the individuals in the discovery cohort were classified into one of the two subgroups according to their SI values. The description of the machine-learning methodology has been provided in detail previously (21). In brief, we applied a cascade of two algorithms, the L_1_-regularized sparse canonical correlation analysis (L_1_-SCCA) (49) and the sparse logistic regression (SLR) analysis (50), thereby effectively reducing the number of parameters (i.e., FCs) to avoid overfitting (35). First, the L_1_-SCCA was applied to the pool of correlation matrices to extract a subset of FCs relevant only to the neural substrates of the SI by eliminating the unwanted effects of NVs. In the present study, the L_1_-SCCA eliminated FCs that were correlated with age, sex, BDI score, and head motion-related parameters that exhibited statistically significant or trend towards differences between the two subgroups (see Table S2). The distributions of these parameters can differ from cohort to cohort, thereby hampering the generalization of the classification scheme (21). We then applied the SLR to further perform dimension reduction to extract the most informative FCs that reflected the neural substrates of the SI. The two algorithms were embedded in a framework of nested cross-validation and LOOCV. In the present study, we used 5-fold CV so that each fold incorporated approximately 24 participants. The output of the SLR is the final set of FCs and the associated weights. The weighted linear summation of the corresponding correlation indices was used to predict an individual’s identity to either of the two subgroups. Namely, *P*(**z**;**w**) = 1 / [1 + exp(–**w**^T^**z**)] determines the identity, where **w** and **z** are the vectors of the normalized correlation indices and the associated weights determined by the SLR, respectively (21). The individual is predicted to belong to SI_L_ or SI_H_ if the corresponding *P* is ≤ or > 0.5, respectively. To evaluate the stability and robustness of the selection of the FCs in the LOOCV, we evaluated the cumulative absolute weight of the form 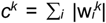 where 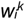 is the weight associated with the *k*-th FC during the *i*-th fold of the LOOCV and the sum runs over all folds. The greater magnitude of *c^k^* indicates a more significant contribution to the classification between the two subgroups. In addition, for each FC, we tested whether the set of weights assigned in the LOOCV was significantly non-zero using a one-sample *t*-test. *P*-values were adjusted for false discovery rate (FDR) based on the Benjamini-Hochberg method (51).

### PET data acquisition and analysis

The participants in the validation cohort underwent a PET scan to evaluate the striatal DA D_2_R availability. The procedures for data acquisition and analysis have been described in detail previously (9, 52). In short, following the intravenous rapid bolus injection of [^11^C]raclopride, a dynamic PET scan was conducted for 60 min. PMOD software (PMOD Technologies Ltd., Zurich, Switzerland) was used to evaluate the temporal radioactivity of [^11^C]raclopride, thereby deriving a parametric image of non-displaceable binding potential (BP_ND_), which represents the spatial distribution of DA D_2_R availability in the brain. Specifically, we estimated BP_ND_ in the left SMST (9) using the three-parameter simplified reference tissue model (53) with the cerebellum as a reference region (37). The boundary of the SMST was determined using the Oxford-GSK-Imanova Striatal Connectivity Atlas (54) which was transformed into the custom template space of the participants. Here, we used Advanced Normalization Tools (ANTs; http://stnava.github.io/ANTs/) to construct this custom template from pairs of PET parametric and T_1_-weighted MR structural images of all the participants. The deformation parameters from a participant’s native to the custom template space were also derived using this procedure. For each participant, the BP_ND_ in the SMST was estimated by calculating the mean of the BP_ND_ within the SMST on the individual’s parametric image in the custom template space.

### Linear regression of DA D_2_R BP_ND_ of [^11^C]raclopride

Using the 15 FCs incorporated in the pSI classifier, we attempted to predict the individual’s striatal DA D_2_R availability. In the LOOCV framework, the individual BP_ND_ of [^11^C]raclopride in the SMST was linearly regressed by the correlation indices of the 15 FCs in the classifier. We incorporated age as an NV in the model, as previous studies have reported age dependence on DA D_2_R properties (55). [Note that sex and BDI were not incorporated as NVs in the model because the validation cohort comprised only male participants, for whom the BDI scores were not available.] Using the derived coefficients (i.e., weights), we predicted the BP_ND_ of the held-out individual as the weighted linear sum of the respective correlation indices. The agreement between the measured and predicted BP_ND_ was evaluated using Pearson’s correlation. To evaluate the reliability of the prediction, we conducted a bootstrapping analysis of 10,000 repetitions, and alternative models were constructed using randomly selected 15 FCs not incorporated in the pSI classifier (i.e., 15 FCs out of 7,488 (= 7503 – 15) FCs). Reliability was evaluated by integrating the cumulative distribution of the pooled correlation indices obtained through the bootstrapping procedure.

## Acknowledgments

This work was supported in part by the Strategic Research Program for Brain Sciences (Integrated Research on Depression, Dementia and Development Disorders) (JP20dm0107094) from the Japan Agency for Medical Research and Development (AMED) to MY and TS, and the Brain/MINDS Beyond (JP21dm0307007 and JP21dm0307008) from the AMED to NY, ERATO (JPMJER1801) supported by Japan Science and Technology Agency (JST) to NY, MEXT Quantum Leap Flagship Program (MEXT Q-LEAP) Grant Number JPMXS0120330644 to NY and MY, and KAKENHI (20H05711) to MY.

