## Supplementary Information for "Neural network of superiority illusion predicts the level of dopamine in striatum"

**Supplementary Information Text**

**Identification of superiority illusion (SI)-related functional connectivities (FCs) for negative trait words based on a low state anxiety population.** Here, we present a supplementary analysis to identify a set of FCs relevant to the SI in the evaluation of negative trait words. Unlike the main analysis, we formed a subset of the discovery cohort by excluding individuals with high state anxiety to the extent allowed by the original dataset. This was intended to attain an improved homogeneity in negative emotional states among the individuals, thereby improving the reliability of the feature selection procedure (see Discussion). We used the state anxiety measurement of the Japanese version of the State-Trait Anxiety Inventory (STAI) available for 91 out of 123 participants in the discovery cohort. The mean and standard deviation (SD) of their state anxiety scores were 37.4 ± 8.4 (range, 20–60). Based on the pre-determined cut-off value (score = 42) (1), a total of 28 participants (31%) were identified as being in a state of high anxiety and were excluded from the subsequent analysis. For stability of the feature selection procedure, the subset further incorporated another 32 participants from the discovery cohort for whom the STAI measurements were not available. We note that this extra group of participants may include some individuals with high state anxiety, so the following results should be considered preliminary findings.

The final subset contained a total of 95 participants (age, 32.4 ± 14.4 yr; 26 females), and the mean and SD of the SI measurements for negative trait words was 0.13 ± 0.21. This mean value was used as a threshold to form two subgroups of participants, $nSI_L$ ($N$ = 69; mean ± SD of nSI = –0.01 ± 0.12) and $nSI_H$ ($N$ = 26; mean ± SD of nSI = 0.33 ± 0.14) below and above the mean nSI (see Fig. S3 and Table S3). We constructed a classifier using the weighted linear summation (WLS) of the respective Pearson correlation indices. The leave-one-out cross-validation procedure revealed that the accuracy of the classification with respect to the actual group identity was 75% (area under the curve [AUC]= 0.78; sensitivity = 91% and specificity = 60%), which was statistically significant (permutation test, $P < 10^{-4}$). The accuracy of classification in the independent validation cohort was 65% (AUC = 0.67; sensitivity = 83%, specificity = 53%). In addition, we applied this classifier to the group with high state anxiety, excluded from the analysis. The accuracy of the classification was 36%, suggesting that individuals with low and high state anxiety could recruit differential neural circuits promoting SI for negative trait words.

**Fig. S1.**

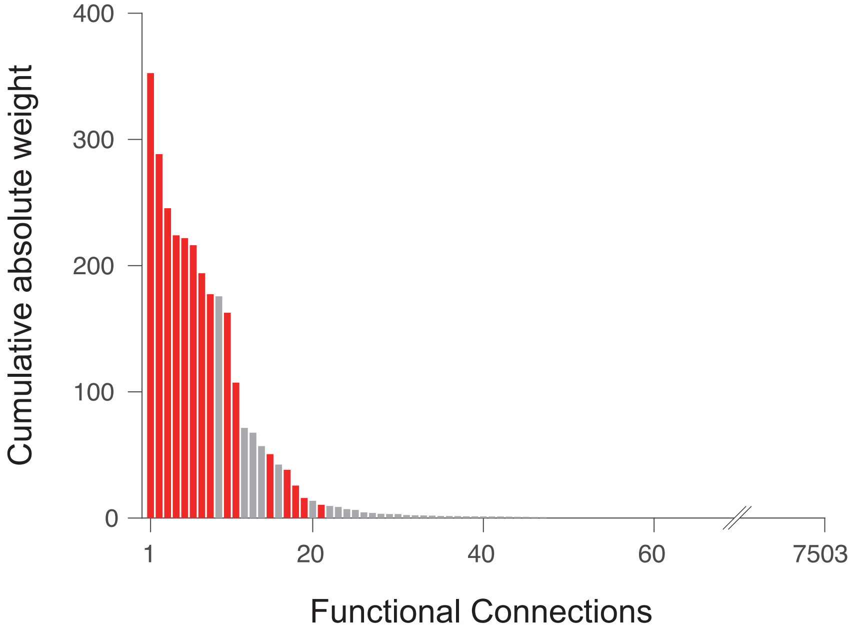


The cumulative absolute weights of functional conectivities (FCs) summed across the leave-one-out cross validation (LOOCV) procedure, indicating the contribution of each FC to the classification of the $pSI_L$ and $pSI_H$ subgroups. A greater cumulative absolute weight represents a larger degree of contribution of the FC to the classifier.

**Fig. S2.**

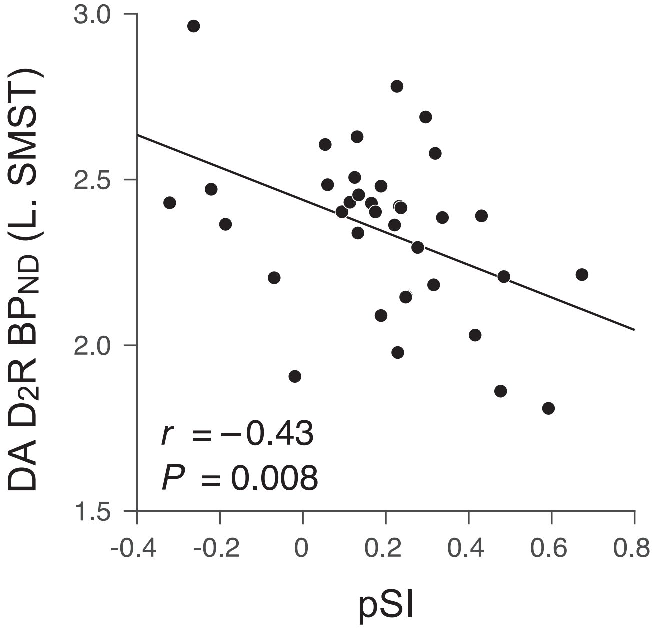


Significant negative correlation between the individual propensity to pSI and the $BP_{ND}$ in the left sensorimotor striatum (SMST) in the validation cohort ($n$ = 36; $r$ = –0.43, $P$ < 0.008).

**Fig. S3.**

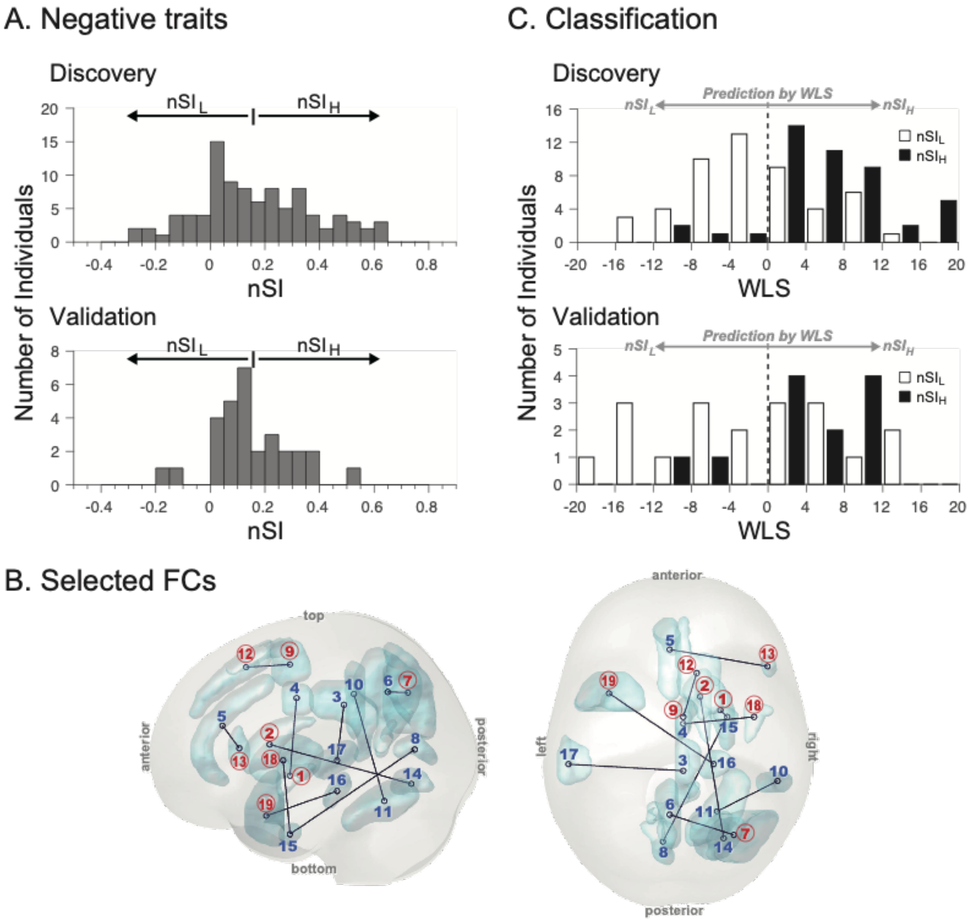


The results of the follow-up analysis for the negative trait words using exclusively the low state-anxiety individuals below the cutoff score of the State-Trait Anxiety Inventory (STAI). (*A*) Distribution of nSI in the discovery and validation samples. (*B*) The 10 functional connectivities (FCs) used in the classification of the $nSI_L$ and $nSI_H$ subgroups as viewed from the left and top of the brain. The ID numbers assigned to the terminal regions correspond to the Region ID in Table S3. The numbers in the circle indicate that the corresponding terminal regions were also present in the 15 FCs selected in the classification of pSI subgroups (see Fig. 2 and Table 2). (C) Distribution of weighted linear summation (WLS) of the 10 FCs in the discovery cohort. The WLS smaller and greater than 0 is classified as $nSI_L$ and $nSI_H$ group, respectively. (*Top*) The number of individuals in the $nSI_L$ (open bar) and $nSI_H$ (filled bar) subgroups in the discovery cohort is shown as a histogram with the WLS width of 4. (*Bottom)* The WLS distribution of the validation cohort.

**Table S1.** Summary of magnetic resonance (MR) imaging parameters. *, the first four volumes were excluded to allow for $T_1$ equilibrium effects.

| | Discovery cohort | Validation cohort |
|---|---|---|
| MRI scanner | Siemens Verio | GE Signa HDx |
| Magnetic field strength (T) | 3.0 | |
| Resting-state fMRI | | |
| Field of view (mm) | 240 | |
| Matrix | 64 × 64 | |
| Number of slices | 33 | 35 |
| Number of volumes* | 204 | |
| In-plane resolution (mm) | 3.75 × 3.75 | |
| Slice thickness (mm) | 3.8 | |
| Repetition time (ms) | 2,000 | |
| Echo time (ms) | 25 | |
| Flip angle (deg) | 90 | |
| Slice acquisition order | Ascending (interleaved) | |
| Instruction to participants | Relax, don't think of anything in particular nor sleep, but keep looking at the crosshair mark in the screen. | |
| Structural scan | | |
| Field of view (mm) | 250 | 256 |
| Matrix | 512 × 512 | 256 × 256 |
| Number of slices | 176 | 196 |
| In-plane resolution (mm) | 0.49 × 0.49 | 1 × 1 |
| Slice thickness (mm) | 1 | |
| Repetition time (ms) | 2,300 | 6.7 |
| Echo time (ms) | 1.9 | |
| Inversion time (ms) | 9 | 12 |

**Table S2.** Head motion-related parameters during the resting-state functional magnetic resonance imaging (rsfMRI) scan in the (*A*) discovery and (*B*) validation cohorts. For parameters involving head rotation (pitch, roll, and yaw), the head radius of 50 mm was assumed. The following four kinds of head motion measures were evaluated: mean relative displacement (MRD) (2), root mean square relative displacement (RMS RD) (3), mean framewise displacement (FD) (4), and mean number of offending frames (NOF) identified by the ArtRepair algorithm (5). In the discovery cohort, parameters exhibiting statistically significant (**P* < 0.05) or trend toward (†*P* < 0.1) differences between the $SI_L$ and $SI_H$ subgroups were incorporated as a nuisance variable in the feature selection (see Materials and Methods).

A. Positive trait words

| | | Discovery | | | Validation | | |
|---|---|---|---|---|---|---|---|
| | | $pSI_L$ | $pSI_H$ | *P* | $pSI_L$ | $pSI_H$ | *P* |
| MRD (mm) | x | 0.013±0.006 | 0.016±0.008 | †0.078 | 0.013±0.006 | 0.010±0.005 | 0.131 |
| | y | 0.061±0.025 | 0.065±0.028 | 0.684 | 0.036±0.018 | 0.031±0.012 | 0.592 |
| | z | 0.052±0.031 | 0.066±0.035 | *0.007 | 0.035±0.021 | 0.027±0.013 | 0.236 |
| | pitch | 0.034±0.012 | 0.038±0.014 | 0.120 | 0.037±0.023 | 0.027±0.009 | 0.390 |
| | roll | 0.013±0.006 | 0.014±0.006 | 0.221 | 0.013±0.004 | 0.012±0.005 | 0.408 |
| | yaw | 0.012±0.007 | 0.014±0.007 | †0.070 | 0.012±0.005 | 0.010±0.004 | 0.277 |
| RMS RD (mm) | x | 0.020±0.010 | 0.023±0.013 | 0.115 | 0.017±0.008 | 0.014±0.007 | 0.123 |
| | y | 0.075±0.031 | 0.079±0.034 | 0.647 | 0.048±0.022 | 0.040±0.015 | 0.277 |
| | z | 0.068±0.035 | 0.090±0.052 | *0.007 | 0.051±0.040 | 0.039±0.019 | 0.372 |
| | pitch | 0.047±0.021 | 0.052±0.022 | 0.189 | 0.057±0.053 | 0.037±0.013 | 0.390 |
| | roll | 0.020±0.012 | 0.020±0.009 | 0.422 | 0.018±0.006 | 0.017±0.008 | 0.733 |
| | yaw | 0.018±0.017 | 0.020±0.011 | 0.112 | 0.016±0.007 | 0.014±0.006 | 0.548 |
| Mean FD (mm) | | 0.16±0.05 | 0.17±0.05 | †0.064 | 0.12±0.07 | 0.09±0.03 | 0.108 |
| Mean NOF (frames) | | 8.30±6.96 | 12.5±9.20 | *0.003 | 6.29±9.47 | 6.50±5.52 | 0.431 |

B. Negative trait words

| | | Discovery | | | Validation | | |
|---|---|---|---|---|---|---|---|
| | | $nSI_L$ | $nSI_H$ | *P* | $nSI_L$ | $nSI_H$ | *P* |
| MRD (mm) | x | 0.014±0.007 | 0.015±0.007 | 0.626 | 0.012±0.006 | 0.010±0.004 | 0.392 |
| | y | 0.062±0.028 | 0.064±0.025 | 0.579 | 0.034±0.017 | 0.032±0.013 | 1.000 |
| | z | 0.052±0.031 | 0.067±0.034 | *0.006 | 0.033±0.019 | 0.027±0.013 | 0.267 |
| | pitch | 0.035±0.013 | 0.037±0.012 | 0.182 | 0.033±0.019 | 0.028±0.014 | 0.392 |
| | roll | 0.014±0.006 | 0.013±0.006 | 0.303 | 0.013±0.005 | 0.011±0.004 | 0.194 |
| | yaw | 0.013±0.007 | 0.013±0.006 | 0.773 | 0.012±0.005 | 0.009±0.003 | 0.229 |
| RMS RD (mm) | x | 0.021±0.011 | 0.022±0.012 | 0.812 | 0.017±0.008 | 0.013±0.006 | 0.217 |
| | y | 0.076±0.034 | 0.078±0.030 | 0.562 | 0.045±0.021 | 0.041±0.015 | 0.635 |
| | z | 0.069±0.036 | 0.091±0.053 | *0.009 | 0.049±0.036 | 0.038±0.019 | 0.267 |
| | pitch | 0.048±0.023 | 0.051±0.020 | 0.306 | 0.051±0.045 | 0.038±0.018 | 0.254 |
| | roll | 0.021±0.012 | 0.018±0.008 | 0.153 | 0.019±0.008 | 0.016±0.006 | 0.281 |
| | yaw | 0.020±0.017 | 0.018±0.011 | 0.576 | 0.017±0.008 | 0.013±0.004 | 0.194 |
| Mean FD (mm) | | 0.16±0.06 | 0.17±0.05 | 0.145 | 0.11±0.06 | 0.10±0.04 | 0.358 |
| Mean NOF (frames) | | 9.63±8.29 | 11.2±8.41 | 0.133 | 7.62±8.40 | 5.06±5.46 | 0.256 |

**Table S3.** Demographic information and the superiority illusion (SI) measurement for negative trait words using the low state anxiety individuals in the discovery and validation cohorts. *P*-values were determined by a Wilcoxon rank sum test for SI, age, and the Beck Depression Inventory (BDI) score, and by a chi-squared test for sex. †The BDI measurements were available only for the discovery cohort.

| Negative trait words | | | | | | | | |
|---|---|---|---|---|---|---|---|---|
| | Discovery cohort (low anxiety individuals) | | | | Validation cohort (low anxiety individuals) | | | |
| | | Subgroup | | | | Subgroup | | |
| Item | All | $nSI_L$ | $nSI_H$ | *P* | All | $nSI_L$ | $nSI_H$ | *P* |
| N | 95 | 69 | 26 | – | 31 | 19 | 12 | – |
| nSI | 0.13±0.21 | -0.01±0.12 | 0.33±0.14 | < 0.001 | 0.11±0.21 | -0.03±0.19 | 0.26±0.10 | < 0.001 |
| Age (y) | 32.4±14.4 | 28.7±10.8 | 36.5±16.7 | 0.03 | 23.6±4.7 | 23.2±3.4 | 24.2±6.3 | 0.87 |
| Sex (M/F) | 50 / 45 | 37 / 32 | 13 / 13 | 0.75 | 31 / 0 | 19 / 0 | 12/ 0 | – |
| BDI† | 4.1±3.6 | 4.7±4.0 | 3.5±3.0 | 0.25 | (n/a) | (n/a) | (n/a) | – |

**Table S4.** Properties of the 10 functional connectivities (FCs) used to classify the $nSI_L$ and $nSI_H$ subgroups formed in the low state-anxiety subset of the discovery cohort. The region name in bold face indicates that it was included also in the set of 15 FCs used to classify the $pSI_L$ and $pSI_H$ subgroups (see Table 2). Contribution was calculated as Weight × | $r(nSI_L) - r(nSI_H)$ |, and the number in parenthesis indicates the relative fraction (%). Lat., laterality; BA, Brodmann area; Contrib., contribution; c., cortex; g., gyrus. *, see Fig. S3.

| FC# | Terminal regions | | | | Mean correlation | | Weight | Contrib. (%) |
|---|---|---|---|---|---|---|---|---|
| | Map ID* | Lat. | Name | BA | $r(nSI_L)$ | $r(nSI_H)$ | | |
| 1 | (6) | L | Precuneous c. | 7 | 0.03 | 0.13 | 8.04 | 0.80 |
| | (7) | **R** | **Lateral occipital c.** (superior) | 31 | | | | (21.3) |
| 2 | (3) | R | Cingulate g. (posterior) | 31 | –0.15 | –0.08 | 9.60 | 0.63 |
| | (17) | L | Superior temporal g. (posterior) | 22 | | | | (16.8) |
| 3 | (11) | **R** | **Cerebellum VI** | – | –0.08 | –0.02 | 6.71 | 0.47 |
| | (10) | **R** | **Supramarginal g.** (posterior) | 40 | | | | (12.5) |
| 4 | (16) | **R** | **Parahippocampal g.** (posterior) | 35 | 0.11 | 0.04 | –5.81 | 0.43 |
| | (19) | L | Temporal pole | 38 | | | | (11.4) |
| 5 | (15) | **R** | **Temporal fusiform c.** (anterior) | 28 | –0.02 | –0.08 | –6.8 | 0.37 |
| | (8) | L | Intracalcarine c. | 18 | | | | (9.8) |
| 6 | (9) | R | Supplementary motor c. | 6 | –0.08 | –0.03 | 6.90 | 0.33 |
| | (12) | R | Superior frontal g. | 6 | | | | (8.8) |
| 7 | (4) | R | Cingulate g. (posterior middle) | 24 | 0.04 | 0.07 | 7.58 | 0.28 |
| | (18) | R | Planum polare | 13 | | | | (7.4) |
| 8 | (5) | L | Paracingulate g. | 32 | 0.05 | 0.01 | –4.49 | 0.22 |
| | (13) | **R** | **Inferior frontal g.** (pars triangularis) | 45 | | | | (5.9) |
| 9 | (2) | **R** | **Caudate** | – | –0.07 | –0.14 | –2.29 | 0.15 |
| | (14) | R | Occipital fusiform g. | 18 | | | | (4.0) |
| 10 | (1) | **R** | **Putamen** | – | 0.00 | 0.03 | 2.53 | 0.08 |
| | (15) | **R** | **Temporal fusiform c.** (anterior) | 28 | | | | (2.1) |
